# An enzyme-based immunodetection assay to quantify SARS-CoV-2 infection

**DOI:** 10.1101/2020.06.14.150862

**Authors:** Carina Conzelmann, Andrea Gilg, Rüdiger Groß, Desirée Schütz, Nico Preising, Ludger Ständker, Bernd Jahrsdörfer, Hubert Schrezenmeier, Konstantin M. J. Sparrer, Thomas Stamminger, Steffen Stenger, Jan Münch, Janis A. Müller

## Abstract

SARS-CoV-2 is a novel pandemic coronavirus that caused a global health and economic crisis. The development of efficient drugs and vaccines against COVID-19 requires detailed knowledge about SARS-CoV-2 biology. Several techniques to detect SARS-CoV-2 infection have been established, mainly based on counting infected cells by staining plaques or foci, or by quantifying the viral genome by PCR. These methods are laborious, time-consuming and expensive and therefore not suitable for a high sample throughput or rapid diagnostics. We here report a novel enzyme-based immunodetection assay that directly quantifies the amount of *de novo* synthesized viral spike protein within fixed and permeabilized cells. This in-cell ELISA enables a rapid and quantitative detection of SARS-CoV-2 infection in microtiter format, regardless of the virus isolate or target cell culture. It follows the established method of performing ELISA assays and does not require expensive instrumentation. Utilization of the in-cell ELISA allows to e.g. determine TCID_50_ of virus stocks, antiviral efficiencies (IC_50_ values) of drugs or neutralizing activity of sera. Thus, the in-cell spike ELISA represents a promising alternative to study SARS-CoV-2 infection and inhibition and may facilitate future research.

**Highlights:**

- Determination of SARS-CoV-2 infection by enzymatically quantifying the expression of viral spike protein in bulk cell cultures
- Targeting a highly conserved region in the S2 subunit of the S protein allows broad detection of several SARS-CoV-2 isolates in different cell lines
- Screening of antivirals in microtiter format and determining the antiviral activity as inhibitory concentrations 50 (IC_50_)

## Introduction

The severe acute respiratory syndrome coronavirus 2 (SARS-CoV-2) emerged as a novel human pathogen at the end of 2019 and spread around the globe within three months. It causes the coronavirus disease 2019 (COVID-19) that if symptomatic manifests as fever, cough, and shortness of breath, and can progress to pneumonia, acute respiratory distress syndrome resulting in septic shock, multi-organ failure and death. As of end of May 2020, more than 376,000 deaths worldwide occurred upon SARS-CoV-2 infection which forced governments to implement strict measures of social distancing to limit the spread of the virus but greatly impacted individual freedom and economy. Due to its high transmissibility, without such harsh interventions its pandemic spread is unlikely to be stopped without the cost of a substantial death toll. Therefore, the development of prophylactics or therapeutics against SARS-CoV-2 is imperative.

SARS-CoV-2 is a positive-sense single-stranded RNA virus with diameters of 60-140 nanometers (Zhu et al., 2020). Like other coronaviruses, SARS-CoV-2 has four structural proteins, the S (spike), E (envelope), M (membrane), and N (nucleocapsid) proteins. The S protein is responsible for allowing the virus to attach to and fuse with the membrane of a host cell. It is primed by the transmembrane serine protease 2 (TMPRSS2) resulting in interactions of the S1 subunit with the angiotensin converting enzyme 2 (ACE2) and rearrangements in S2 to form a six-helix bundle structure that triggers fusion of the viral with the cellular membrane (Hoffmann et al., 2020; Q. Wang et al., 2020; Xia et al., 2020a, 2020b). Compounds interfering with the binding of the S protein to ACE2 (Ou et al., 2020; C. Wang et al., 2020) or inhibiting TMPRSS2 or formation of the six-helix bundle also suppress infection by SARS-CoV-2 (Hoffmann et al., 2020; Xia et al., 2020a, 2020b). SARS-CoV-2 is now intensely investigated to understand viral biology and pathogenesis and to develop antiviral drugs and vaccines. Techniques to study and quantify SARS-CoV-2 infection and replication in cell culture have quickly evolved in the past months, partially inspired by methods developed for the related SARS-CoV or other (corona-) viruses.

SARS-CoV-2 infection is mainly quantified by determining the number of infectious particles by counting virus-induced plaques or foci after staining with crystal violet, neutral red (Keil et al., 2020; Ma et al., 2020; Runfeng et al., 2020; Xia et al., 2020a) or specific antibodies for SARS-CoV-2 antigens, e.g. against the N protein (Chu et al., 2020; X. Liu et al., 2020). These antibodies are either labelled directly with horseradish peroxidase (HRP) or fluorophores, or are detected by a corresponding secondary labelled antibody. The number of infected cells is then detected by immunofluorescence microscopy, flow cytometry, or by manual counting by microscope or with the help of computational algorithms (J. Liu et al., 2020; Ma et al., 2020; Ou et al., 2020; Runfeng et al., 2020). Other methods to quantify infection rates are detection of cell-associated RNA by RT-qPCR (Monteil et al., 2020; Runfeng et al., 2020) or viral proteins by western blotting (Ou et al., 2020), which are expensive and unsuitable for large sample numbers. Alternatively, with prolonged waiting times until results are available, viral replication may be measured by determining the RNA or infectious titers of progeny virus released from infected cells by RT-qPCR (Chu et al., 2020; J. Liu et al., 2020; Yao et al., 2020), or tissue culture infectious dose 50 (TCID_50_) endpoint titrations (Chin et al., 2020; Manenti et al., 2020) and plaque assays (Keil et al., 2020; Ma et al., 2020; Runfeng et al., 2020; Xia et al., 2020a), respectively. All these assays are well established and validated but have the downside of being laborious, time-consuming, lacking specificity, and the difficulty to increase sample sizes to perform analysis in microtiter format which is substantial in the search for antivirals or in diagnostics. Instead of counting infected cells or quantifying RNA, we here developed an in-cell ELISA that directly quantifies SARS-CoV-2 infection by detecting newly synthesized S protein. The assay allows detection of all SARS-CoV-2 isolates tested and can be easily performed in any format including 96-well plates. It can be used to measure the TCID_50_, to screen for antivirals, and to determine antiviral potencies of drugs (as inhibitory concentration 50), neutralizing sera or antibodies in a timely and cost-effective manner, within only two days.

## Materials and Methods

### Cell culture

Vero E6 (*Cercopithecus aethiops* derived epithelial kidney) cells were grown in Dulbecco’s modified Eagle’s medium (DMEM, Gibco) which was supplemented with 2.5% heat-inactivated fetal calf serum (FCS), 100 units/ml penicillin, 100 μg/ml streptomycin, 2 mM L-glutamine, 1 mM sodium pyruvate, and 1x non-essential amino acids. Caco-2 (human epithelial colorectal adenocarcinoma) cells were grown in the same media but with supplementation of 10% FCS. Calu-3 (human epithelial lung adenocarcinoma) cells were cultured in Minimum Essential Medium Eagle (MEM, Sigma #M4655) supplemented with 10% FCS, 100 units/ml penicillin, 100 μg/ml streptomycin, 1 mM sodium pyruvate, and 1x non-essential amino acids. All cells were grown at 37°C in a 5% CO_2_ humidified incubator.

### Virus strains and virus propagation

Viral isolate BetaCoV/France/IDF0372/2020 (#014V-03890) and BetaCoV/Netherlands/01/NL/2020 (#010V-03903) were obtained through the European Virus Archive global. Virus was propagated by inoculation of 70% confluent Vero E6 in 75 cm^2^ cell culture flasks with 100 μl SARS-CoV-2 isolates in 3.5 ml serum-free medium containing 1 μg/ml trypsin. Cells were incubated for 2 h at 37°C, before adding 20 ml medium containing 15 mM HEPES. Cells were incubated at 37°C and supernatant harvested at day 3 post inoculation when a strong cytopathic effect (CPE) was visible. Supernatants were centrifuged for 5 min at 1,000 × g to remove cellular debris, and then aliquoted and stored at −80°C as virus stocks. Infectious virus titer was determined as plaque forming units or TCID_50_.

### Virus isolation from patient samples

To isolate SARS-CoV-2 from patient samples, 50,000 Vero E6 cells were seeded in 24-well plates in 500 μl medium incubated over night at 37°C. The next day, medium was replaced by 400 μl of 2.5 μg/ml amphotericin B containing medium. Then, 100 μl of throat swabs that were tested positive for SARS-CoV-2 by qRT-PCR were titrated 5-fold on the cells and incubated for 3 to 5 days. Upon visible CPE, supernatant was taken and virus expanded by inoculation of Vero E6 cell in 75 cm2 flasks and propagated as above described, resulting in the two viral isolates BetaCoV/Germany/Ulm/01/2020 and BetaCoV/Germany/Ulm/02/2020.

### Plaque assay

To determine plaque forming units (PFU), SARS-CoV-2 stocks were serially diluted 10-fold and used to inoculate Vero E6 cells. To this end, 800,000 Vero E6 cells were seeded per 12 well in 1 ml medium and cultured overnight to result in a 100% confluent cell monolayer. Medium was removed, cells were washed once with PBS and 400 μl PBS were added. Cells were then inoculated with 100 μl of titrated SARS-CoV-2 and incubated for 1 to 3 h at 37°C with shaking every 15 to 30 min. Next, cells were overlayed with 1.5 ml of 0.8% Avicel RC-581 (FMC Corporation) in medium and incubated for 3 days. Cells were fixed by adding 1 ml 8% paraformaldehyde (PFA) and incubation at room temperature for 45 min. Supernatant was discarded, cells were washed with PBS once, and 0.5 ml of staining solution (0.5% crystal violet and 0.1% triton in water) was added. After 20 min incubation at room temperature, the staining solution was washed off with water, virus-induced plaques were counted, and PFU per ml calculated. Based on the applied PFU per cell the MOIs were calculated.

### TCID_50_ endpoint titration

To determine the tissue culture infectious dose 50 (TCID_50_), SARS-CoV-2 stocks were serially diluted 10-fold and used to inoculate Vero E6 or Caco-2 cells. To this end, 6,000 Vero E6 or 10,000 Caco-2 cells were seeded per well in 96 flat bottom well plates in 100 μl medium and incubated over night before 62 μl fresh medium was added. Next, 18 μl of titrated SARS-CoV-2 of each dilution was used for inoculation, resulting in final SARS-CoV-2 dilutions of 1:10^1^ to 1:10^9^ on the cells in sextuplicates. Cells were then incubated for 5 days and monitored for CPE. TCID_50_/ml was calculated according to Reed and Muench.

### Establishment of the in-cell SARS-CoV-2 ELISA

To establish detection of SARS-CoV-2 infection, 6,000 Vero E6 or 10,000 Caco-2 target cells were seeded in 96 well plates in 100 μl. The next day, 62 μl fresh medium was added and the cells were inoculated with 18 μl of a 10-fold titration series of SARS-CoV-2. One to three days later, SARS-CoV-2 S protein staining was assessed using an anti-SARS-CoV-2 S protein antibody. To this end, cells were fixed by adding 180 μl 8% PFA and 30 min of room temperature incubation. Medium was then discarded and the cells permeabilized for 5 min at room temperature by adding 100 μl of 0.1% Triton in PBS. Cells were then washed with PBS and stained with 1:1,000, 1:5,000 or 1:10,000 diluted mouse anti-SARS-CoV-2 S protein antibody 1A9 (Biozol GTX-GTX632604) in antibody buffer (PBS containing 10% (v/v) FCS and 0.3% (v/v) Tween 20) at 37°C. After one hour, the cells were washed three times with washing buffer (0.3% (v/v) Tween 20 in PBS) before a secondary anti-mouse or anti-rabbit antibody conjugated with HRP was added (1:10,000, 1:15,000, 1:20,000 or 1:30,000) and incubated for 1 h at 37°C. Following four times of washing, the 3,3’,5,5’-tetramethylbenzidine (TMB) peroxidase substrate (Medac #52-00-04) was added. After 5 min light-protected incubation at room temperature, reaction was stopped using 0.5 M H2SO4. The optical density (OD) was recorded at 450 nm and baseline corrected for 620 nm using the Asys Expert 96 UV microplate reader (Biochrom).

### SARS-CoV-2 infection and inhibition assay

*This is the final protocol established during this study and applied to analyze SARS-CoV-2 infection and inhibition*. To determine SARS-CoV-2 infection, 12,000 Vero E6 or 30,000 Caco-2 target cells were seeded in 96 well plates in 100 μl. The next day, fresh medium and the respective compound of interest (chloroquine (Sigma-Aldrich #C6628); lopinavir (Selleck Chemicals #S1380); EK1 (Core Facility Functional Peptidomics, Ulm); remdesivir (Selleck Chemicals #S8932)) was added and the cells inoculated with the desired multiplicity of infection (MOI; based on PFU per cell) of SARS-CoV-2 in a total volume of 180 μl. Alternatively, virus was preincubated with the compound (human (Sigma-Aldrich #L8402) or chicken (Sigma-Aldrich #L4919) lysozyme) and the mix used for inoculation. Two days later, infection was quantified by detecting SARS-CoV-2 S protein. To this end, cells were fixed by adding 180 μl 8% PFA and 30 min of room temperature incubation. Medium was then discarded and cells permeabilized for 5 min at room temperature by adding 100 μl of 0.1% Triton in PBS. Cells were then washed with PBS and stained with 1:5,000 diluted mouse anti-SARS-CoV-2 S protein antibody 1A9 (Biozol GTX-GTX632604) in antibody buffer (PBS containing 10% (v/v) FCS and 0.3% (v/v) Tween 20) at 37°C. After one hour, the cells were washed three times with washing buffer (0.3% (v/v) Tween 20 in PBS) before a secondary anti-mouse antibody conjugated with HRP (Thermo Fisher #A16066) was added (1:15,000) and incubated for 1 h at 37°C. Following four times of washing, the TMB peroxidase substrate (Medac #52-00-04) was added. After 5 min light-protected incubation at room temperature, reaction was stopped using 0.5 M H2SO4. The optical density (OD) was recorded at 450 nm and baseline corrected for 620 nm using the Asys Expert 96 UV microplate reader (Biochrom). Values were corrected for the background signal derived from uninfected cells and untreated controls were set to 100% infection.

### SARS-CoV-2 neutralization assay

Sera was obtained before the SARS-CoV-2 outbreak or from convalescent COVID-19 patients (confirmed by symptoms and positive SARS-CoV-2 RT-qPCR from nasopharyngeal swabs) tested for seroconversion by IgG/IgA ELISA (Euroimmun #EI 2606-9601 G/ #EI 2606-9601 A) according to the manufacturers’ instructions and IgG/IgM chemiluminescent immunoassay (Shenzhen New Industries Biomedical Engineering, #130219015M/ #130219016M) performed fully-automated in a Maglumi 800. To quantify neutralizing activity of the sera, 30,000 Caco-2 target cells were seeded in 96 well plates in 100 μl and the next day 62 μl fresh medium was added. The sera were heat-inactivated (30 min at 56°C), titrated 2-fold starting with a 5-fold dilution, and mixed 1:1 with SARS-CoV-2 France/IDF0372/2020. After 90 min incubation at room temperature, the mix was used to infect the cells with 18 μl in triplicates at a MOI of 0.01. Two days later, SARS-CoV-2 S protein expression was quantified as described above.

### TCID_50_ determination by in-cell SARS-CoV-2 ELISA

Vero E6 cells were inoculated as described above for the TCID_50_ endpoint titration. Cells were then incubated and CPE development observed by microscopy. At day 4 cells were then fixed (8% PFA), permeabilized (0.1% Triton), stained (1:5,000 1A9; 1:15,000 anti-mouse-HRP), visualized (TMB) and detected in a microplate reader as described above. Infected wells were defined as having a higher signal than the uninfected control plus three times the standard deviation. TCID_50_/ml was calculated as described.

### Cell viability assay

The effect of investigated compounds on the metabolic activity of the cells was analyzed using the CellTiter-Glo^®^ Luminescent Cell Viability Assay (Promega #G7571). Metabolic activity was examined under conditions corresponding to the respective infection assays. The CellTiter-Glo^®^ assay was performed according to the manufacturer’s instructions. Briefly, medium was removed from the culture after 2 days of incubation and 50% substrate reagent in PBS was added. After 10 min, the supernatant was transferred into white microtiter plates and luminescence measured in an Orion II Microplate Luminometer (Titertek Berthold). Untreated controls were set to 100% viability.

### Peptide synthesis

EK1 (Xia et al., 2020b, 2020a) was synthesized automatically on a 0.10 mmol scale using standard Fmoc solid phase peptide synthesis techniques with the microwave synthesizer (Liberty blue; CEM). Briefly, Fmoc protecting groups were removed with 20% piperidine in N,N-dimethylformamide (DMF) and amino acid were added in 0.2 molar equivalent together with a 0.5 molar equivalent of O-benzotriazole-N,N,N’N’-tetramethyluronium-hexafluoro-phosphate and a 2 molar equivalent of diisopropylethylamine. The coupling reaction was performed with microwaves in a few minutes followed by a DMF wash. Once the synthesis was completed, the peptide was cleaved in 95% trifluoroacetic acid, 2.5% triisopropylsilane, and 2.5% H2O for one hour. The peptide residue was precipitated and washed with cold diethyl ether and allowed to dry under vacuum to remove residual ether. The peptide was purified using reversed phase preparative high-performance liquid chromatography (HPLC; Waters) in an acetonitrile/water gradient under acidic conditions on a Phenomenex C18 Luna column (5 mm pore size, 100 Å particle size, 250 - 21.2 mm). Following purification, the peptide was lyophilized on a freeze dryer (Labconco) for storage prior to use. The purified peptide mass was verified by liquid chromatography mass spectroscopy (LCMS; Waters).

### Statistical analysis

The determination of the inhibitory concentration 50 (IC_50_) or inhibitory titer 50 by four-parametric nonlinear regression and one-way ANOVA followed by Bonferroni’s multiple comparison test (ns not significant, * P < 0.01, ** P < 0.001, *** P < 0.0001) were performed using GraphPad Prism version 8.2.1 for Windows, GraphPad Software, San Diego, California USA, www.graphpad.com.

## Results

Inspired by the well-established Zika virus infection assay that quantifies the viral envelope (E) protein by a horseradish peroxidase (HRP) coupled antibody (Aubry et al., 2016; Conzelmann et al., 2019; Müller et al., 2018, 2017; Röcker et al., 2018) we here aimed to detect the spike (S) protein of SARS-CoV-2 by a similar approach. To adapt this in-cell ELISA to measure SARS-CoV-2 infection, we made use of the anti-SARS-CoV/SARS-CoV-2 antibody 1A9 that targets the highly conserved loop region between the HR1 and HR2 in the S2 subunit of the S protein (Ng et al., 2014; Walls et al., 2020) (nextstrain.org (Hadfield et al., 2018)). Two SARS-CoV-2 permissive cell lines, Vero E6 (African green monkey epithelial kidney cells) and Caco-2 (heterogeneous human epithelial colorectal adenocarcinoma cells), were seeded in 96-well plates and inoculated with increasing multiplicities of infection (MOIs) of a SARS-CoV-2 isolate from France (BetaCoV/France/IDF0372/2020). After 2, 24, 48 or 72 hours, cells were fixed, permeabilized, and stained with 1:1,000, 1:5,000, or 1:10,000 dilutions of the anti-SARS-CoV-2 S protein antibody 1A9 for 1 hour. After washing, a 1:20,000 dilution of a secondary HRP-coupled anti-mouse antibody was added, cells were incubated for 1 hour, washed again before TMB peroxidase substrate was added. After 5 min, reaction was stopped using H2SO4 and optical density (OD) recorded at 450 nm and baseline corrected for 620 nm using a microplate reader (Biochrom).

Already at day 1, we observed a significant increase in ODs upon infection with the highest MOI in Vero E6 (Fig. 1a) and Caco-2 (Fig. 1b) cells. At day 2, even the lowest MOI of 0.005 resulted in an OD signal over background in Vero E6 cells, and a maximum OD of 0.157 ± 0.004 after infection with a MOI of 0.05. Higher MOIs resulted in reduced ODs in both cell lines because of virus-induced cytopathic effect (CPE) resulting in detached cells, as monitored by light microscopy. S protein present in the viral inoculum did not result in a significantly increased OD as compared to uninfected controls, as shown in the controls experiments (day 0) in both cell lines. Thus, using a combination of a S protein-specific antibody and a secondary detection antibody allows to detect SARS-CoV-2 infected cells by in-cell ELISA, with readily detectable ODs already 2 days post infection.

**Fig. 1.**
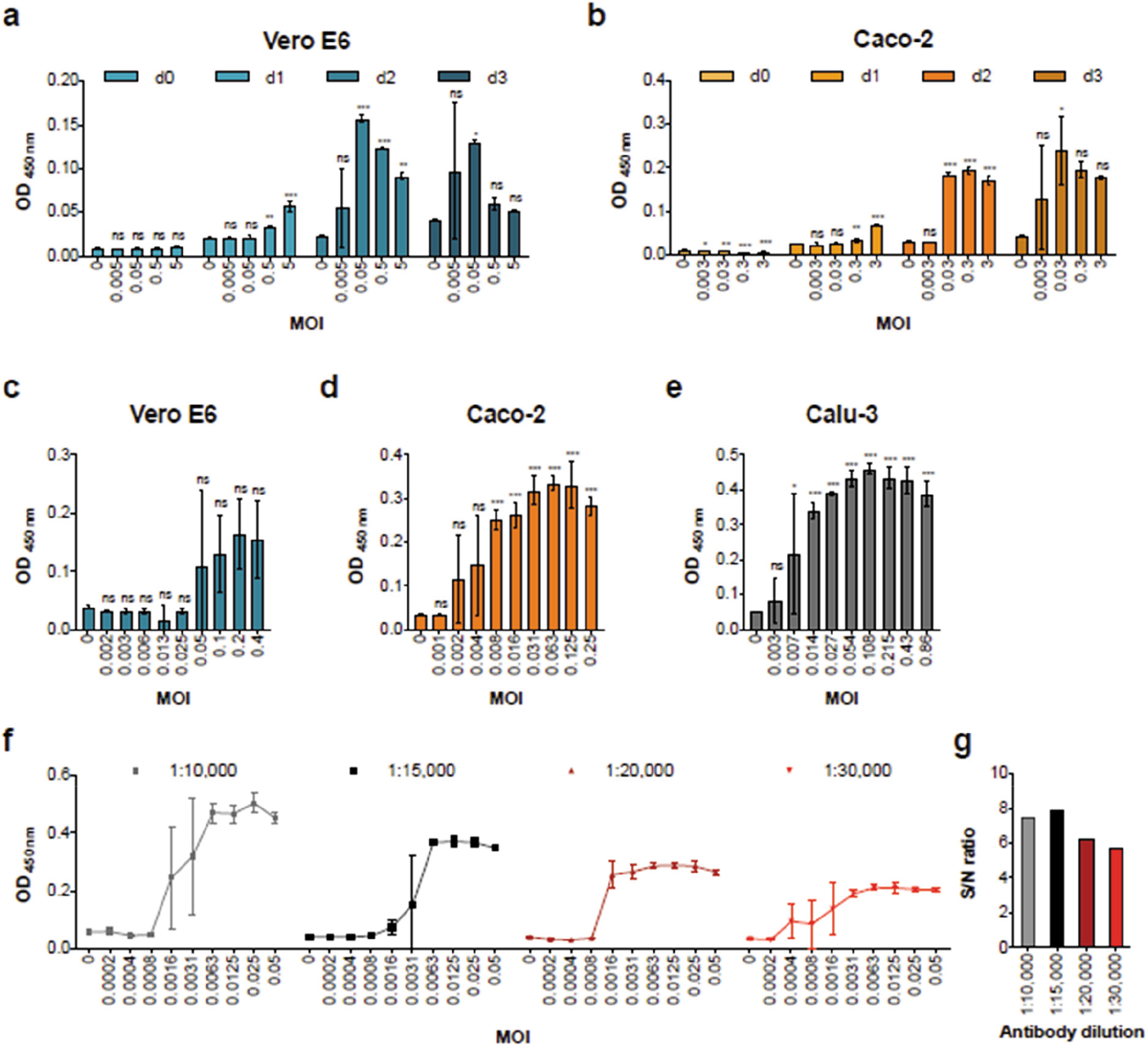
Establishment of an in-cell S protein ELISA to quantify SARS-CoV-2 infection. **a, b)** Time course of S protein expression in infected Vero E6 and Caco-2 cells as detected by in-cell ELISA. Vero E6 (a) and Caco-2 (b) cells were inoculated with increasing MOIs of a SARS-CoV-2 isolate from France. In-cell ELISA (1:5,000 (10 ng/well) 1A9 antibody; 1:20,000 (2.5 ng/well) HRP-antibody) was performed after 2 hours (d0) or 1, 2 or 3 days post infection. **c, d, e)** ELISA signal correlates with viral input dose. Vero E6 (c), Caco-2 (d), or Calu-3 (e) cells were inoculated with serial two-fold dilutions of SARS-CoV-2 and infections rates were determined 2 days later by in-cell ELISA. **f)** Titration of secondary antibody to optimize assay sensitivity applying 5 (1:10,000), 3.3 (1:15,000), 2.5 (1:20,000) or 1.7 ng/well (1:30,000). Caco-2 cells infected with indicated MOIs of SARS-CoV-2 and stained 2 days later with anti-S protein antibody were treated with four dilutions of the HRP-coupled secondary antibody before OD was determined. **g)** Corresponding maximum signal-to-noise (S/N) ratios observed in Fig. 1f. All values show in panels a-e are means of raw data obtained from technical triplicates ± sd. ns not significant, * P < 0.01, ** P < 0.001, *** P < 0.0001 (by one-way ANOVA with Bonferroni’s post-test).

We next inoculated both cell lines and the SARS-CoV-2 susceptible lung cell line Calu-3 (human epithelial lung adenocarcinoma cells) with serial 2-fold dilutions of SARS-CoV-2 and performed the in-cell ELISA 2 days later. A viral inoculum dependent increase in the ODs was detected in Vero E6 cells after infection with a MOI ≥ 0.05 (Fig. 1c), in Caco-2 cells already highly significant with a MOI of ≥ 0.008 (Fig. 1d) and in Calu-3 cells at a MOI of 0.014 (Fig. 1e). Thus, under these experimental conditions, Caco-2 and Calu-3 cells allow a more sensitive detection of SARS-CoV-2 infection and replication as Vero E6 cells.

To optimize assay sensitivity, i.e. the signal-to-noise (S/N) ratio, we evaluated different secondary antibody dilutions. For this, Caco-2 cells were inoculated with SARS-CoV-2 (MOIs of 0.0002 to 0.05), fixed at day 2, and stained with the anti-S protein antibody. Thereafter, four different dilutions of the HRP-coupled secondary antibody were added. OD measurements revealed that highest ODs were obtained with 10,000-fold diluted secondary antibody (Fig. 1f). However, when calculating the S/N ratios (OD of infected wells divided by OD of uninfected cells), also the 15,000-fold dilution revealed a similar assay sensitivity with maximum S/N values of 7.9 as compared to 7.5 for the 1:10,000 dilution (Fig. 1g). Thus, all subsequent experiments were performed in Caco-2 cells that were seeded at a density of 30,000 cells per well to increase ODs, and stained with 5,000-fold diluted anti-S and 15,000-fold diluted secondary antibodies.

Having demonstrated that the in-cell ELISA quantifies infection by a French SARS-CoV-2 isolate, we wanted to validate that isolates from other geographic areas are also detected. The French isolate clusters with the reference Wuhan-Hu-1/2019 isolate whereas the Netherlands/01 strain can be grouped to clade A2a (nextstrain.org (Hadfield et al., 2018)). The antibody-targeted S2 domain is generally conserved between SARS-CoV-2 strains which should allow detection (Ng et al., 2014; Walls et al., 2020) (nextstrain.org (Hadfield et al., 2018)). To test this, Caco-2 cells were inoculated with increasing MOIs of the Netherlands/01 isolate as well as two isolates from Ulm, Southern Germany. Intracellular S protein expression was determined 2 days later by in-cell ELISA. As shown in Fig. 2, virus infection was readily detectable even upon infection with very low MOIs, suggesting that the ELISA may be applied to all SARS-CoV-2 isolates.

**Fig. 2.**
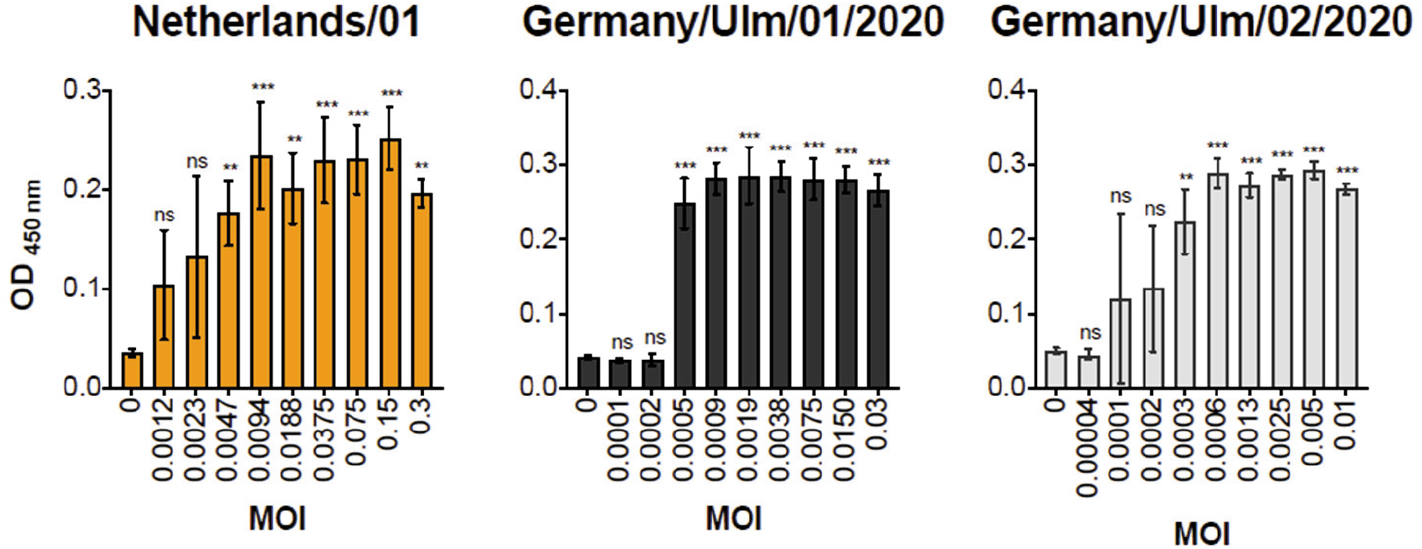
The in-cell S protein ELISA detects SARS-CoV-2 isolates from different geographic regions. Caco-2 cells were infected with increasing MOIs of three SARS-CoV-2 isolates and intracellular S protein expression was quantified 2 days later by in-cell ELISA. Data shown represent means of raw data obtained from technical triplicates ± sd. ns not significant, ** P < 0.001, *** P < 0.0001 (by one-way ANOVA with Bonferroni’s post-test).

Results shown in Fig. 2 indicate that the assay allows to detect infected wells even after inoculation with very low viral MOIs, e.g. a MOI of 0.0003 of the Ulm/01/2020 isolate resulted in a significantly increased OD as compared to uninfected controls. We were wondering whether this high sensitivity and ease of quantitation may also allow to determine the TCID_50_ of virus stocks, that is usually done on Vero E6 cells by manually counting infected wells using a microscope. To test this, we titrated virus, inoculated Vero E6 cells and incubated them for 4 days. We identified infected wells by eye (Fig. 3a), but also performed the in-cell ELISA and set a threshold of three times the standard deviation above the uninfected control to determine the number of infected wells per virus dilution (Fig. 3b). The subsequent calculation of TCID_50_/ml by Reed and Muench revealed exactly the same viral titer for the in-cell ELISA (Fig. 3b) as for microscopic evaluation (Fig. 3a) showing that the established ELISA is suitable for determination of viral titers.

**Fig. 3.**
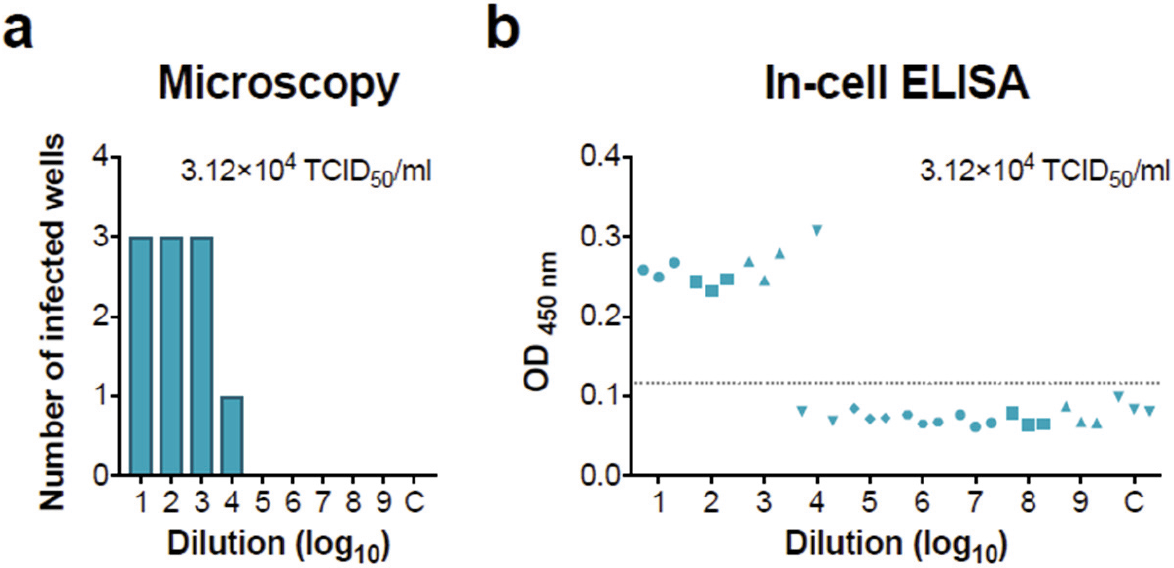
Utilization of the in-cell ELISA to determine the TCID_50_ of SARS-CoV-2 stocks. A stock of the French SARS-CoV-2 isolate was titrated 10-fold and used to inoculate Vero E6 cells in triplicates. At day 4 post infection, the number of infected wells was determined by **a)** microscopically evaluating the CPE or **b)** performing the SARS-CoV-2 S protein in-cell ELISA. Grey line illustrates the threshold of 0.117 (three times the sd added to the uninfected control) used to determine infected wells. The corresponding titer determined according to Reed and Muench is shown as inlet in both figures.

**Fig. 4.**
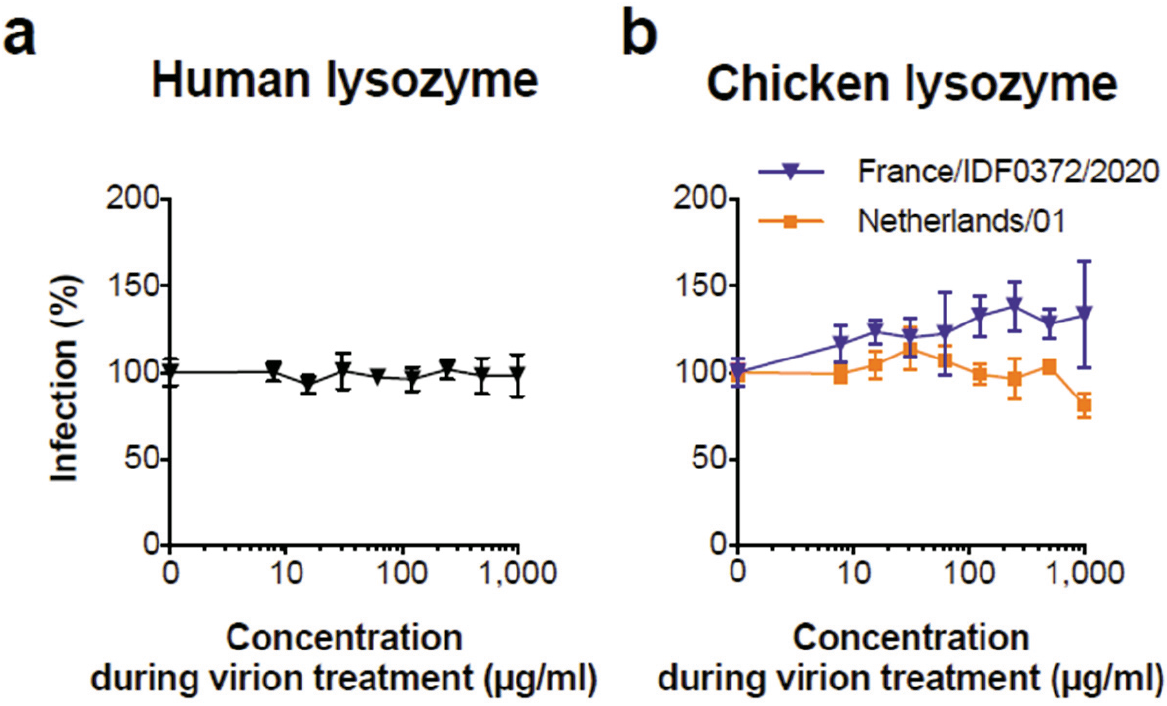
Effect of lysozyme on SARS-CoV-2 infection. **a)** Human lysozyme was incubated with a French SARS-CoV-2 isolate and **b)** chicken lysozyme with a French and Dutch isolate for 2 hours at 37°C before these mixtures were used to infect Caco-2 cells. In-cell ELISA was performed at day 2 post infection. Uninfected controls were subtracted and data normalized to infection rates in absence of lysozyme. Values represent means of 3 technical replicates ± sd.

We next set out to analyse whether lysozyme, a well-known antimicrobial enzyme that is abundant in body fluids such as tears (McDermott, 2013), saliva (Petit and Jollès, 1963), human milk (Andreas et al., 2015; Chanan et al., 1964; Koenig et al., 2005) and mucus (Dajani et al., 2005) may affect SARS-CoV-2 infection. To this end, the French viral isolate was treated with lysozyme purified from human neutrophils, and then used to infect Caco-2 cells. Simultaneously, two more SARS-CoV-2 isolates were treated with lysozyme from chicken egg white and inoculated on Caco-2 cells. In-cell S protein ELISA performed 2 days later demonstrated that none of the lysozyme preparations inhibited viral infection, suggesting that this innate immune defence enzyme does not protect against SARS-CoV-2 infection in saliva or mucus of the respiratory tract.

We then examined whether the ELISA allows to determine the antiviral activity of known SARS-CoV-2 inhibitors. Caco-2 cells were treated with serial dilutions of the small SARS-CoV-2 inhibiting molecules chloroquine (Jeon et al., 2020; M. Wang et al., 2020), lopinavir (Jeon et al., 2020), remdesivir (Jeon et al., 2020; M. Wang et al., 2020) and the peptide inhibitor EK1 (Xia et al., 2020b, 2020a), and were then infected with SARS-CoV-2. In-cell ELISAs performed 2 days later demonstrated a concentration-dependent antiviral activity of the tested compounds reflecting typical dose-response curves of antiviral agents (Fig. 5a-d). This also allowed the calculation of the inhibitory concentration 50 (IC_50_) values, i.e. 23.9 μM for chloroquine (Fig. 5a), 21.0 μM for lopinavir (Fig. 5b), 32.4 nM for remdesivir (Fig. 5c), and 303.5 nM for EK1 (Fig. 5d). These values are in the same range as previously reported on Vero E6 cells (1.13 - 7.36 μM for chloroquine (Jeon et al., 2020; J. Liu et al., 2020; M. Wang et al., 2020), 9.12 μM for lopinavir (Jeon et al., 2020), 770 nM for remdesivir (M. Wang et al., 2020), and 2,468 nM for EK1 (Xia et al., 2020a)), and demonstrate that the in-cell S protein ELISA can be easily adapted to determine antiviral activities of candidate drugs. Cytotoxicity assays that were performed simultaneously in the absence of virus revealed no effects on cell viability by antivirally active concentrations of lopinavir, remdesivir and EK1 (Fig. 5b-d). However, reduced cellular viability rates were observed in the presence of chloroquine concentrations >1 μM (Fig. 5a), which is in line with the fact that part of the anti-SARS-CoV-2 activity of this anti-malaria drug is attributed to its interference with cell organelle function (J. Liu et al., 2020; Mauthe et al., 2018).

**Fig. 5.**
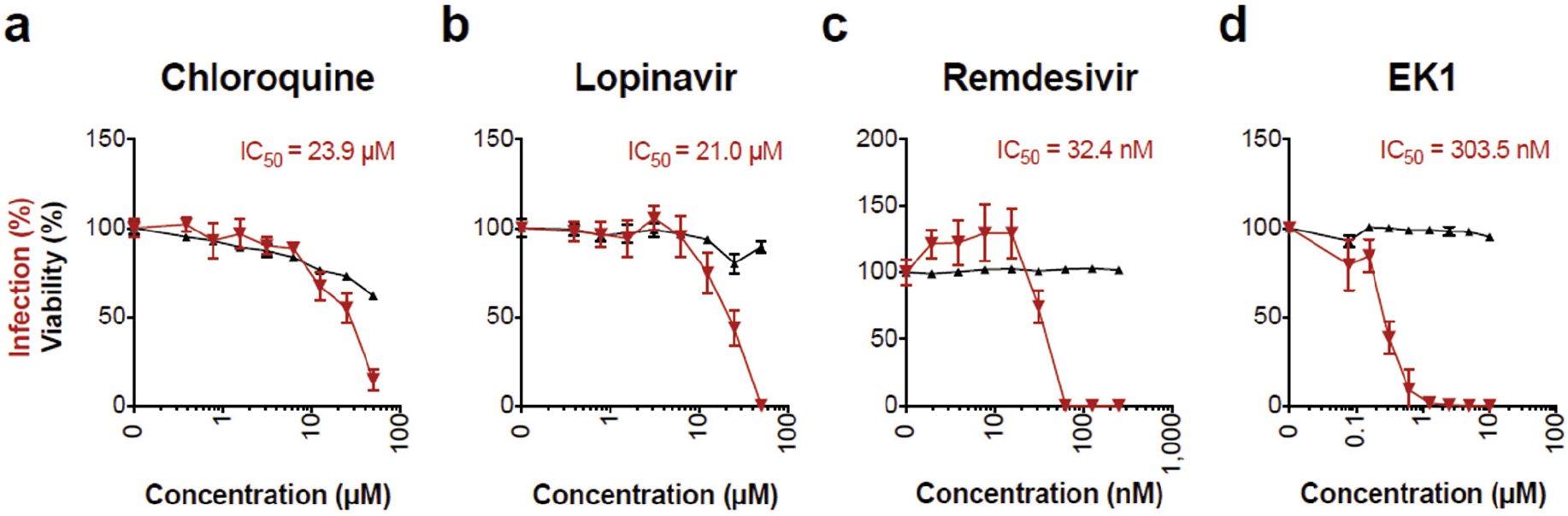
Inhibition of SARS-CoV-2 infection by antivirals. **a-d)** Caco-2 cell treated with chloroquine (a), lopinavir (b), remdesivir (c) or EK1 peptide (d) were infected with SARS-CoV-2 and infection rates were determined 2 days later by in-cell S protein ELISA. Uninfected controls were subtracted and values normalized to infection rates in absence of compound. Shown are means of 4 biological replicates ± sem (chloroquine, lopinavir) or 3 technical replicates ± sd (remdesivir, EK1). Cell viability of Caco-2 cells treated for 2 days with indicated concentrations of drugs was analysed by CellTiter-Glo^®^ Glo assay. Values shown are means of 3 technical replicates ± sd. Inhibitory concentrations 50 (IC_50_) were calculated by nonlinear regression.

Finally, we evaluated whether the assay determines the neutralization activity of serum from SARS-CoV-2 convalescent individuals. For this, sera that were tested positive or negative for anti-SARS-CoV-2 immunoglobulins, were serially titrated and incubated with SARS-CoV-2 for 90 minutes at room temperature before inoculation of Caco-2 cells. Two days later, we performed the in-cell ELISA as described. As shown in Fig. 6, the two control sera, that were obtained before the COVID-19 outbreak or shown to contain no SARS-CoV-2 immunoglobulins, did not affect infection. In contrast, both COVID-19 sera neutralized SARS-CoV-2 infection (Fig. 6). Serum 1 resulted in a more than 50% inhibition at a titer of 640 and Serum 2 already neutralized SARS-CoV-2 at the 1,280-fold dilution. This confirms that the in-cell ELISA is suitable to detect neutralizing sera. Furthermore, analogous to the IC_50_, we calculated the “inhibitory titers 50” using nonlinear regression, and determined titers of 654 and 1,076 respectively. These titers corresponded well to the presence of immunoglobulins which suggests that the here established method can be used to detect and quantify the neutralizing capacities of sera from COVID-19 patients.

**Fig. 6.**
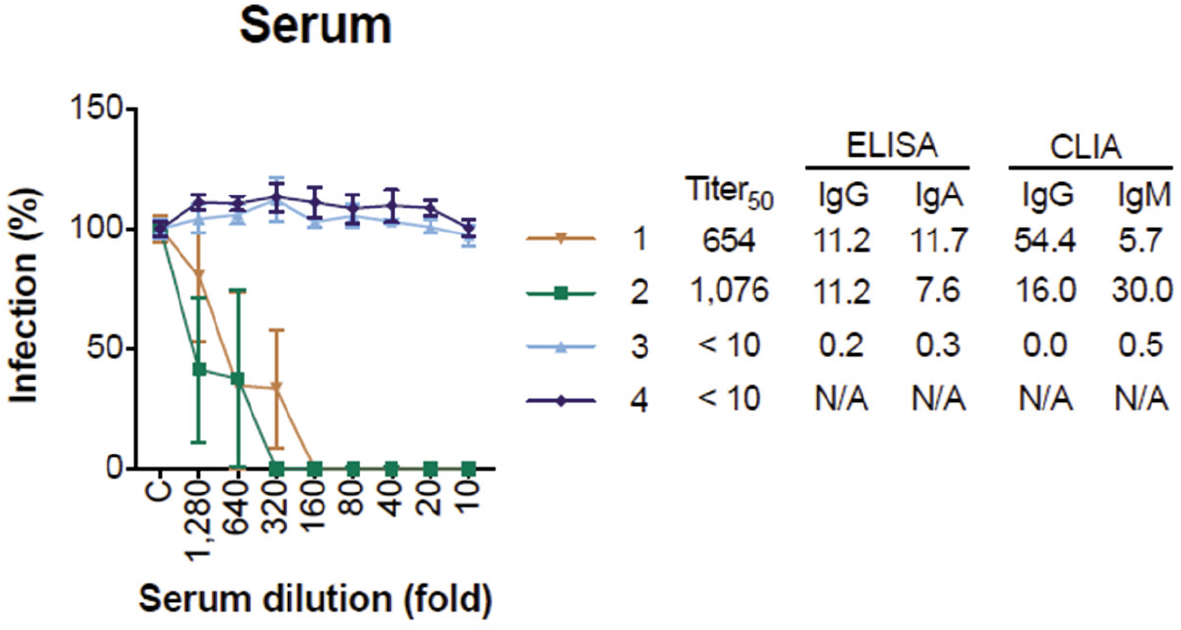
Adaption of the in-cell spike ELISA to determine SARS-CoV-2 neutralizing titers of sera. Sera from convalescent COVID-19 patient or control sera were incubated with a French SARS-CoV-2 isolate for 90 minutes at room temperature and the mixtures were used to infect Caco-2 cells. In-cell ELISA was performed at day 2 post infection. Uninfected controls were subtracted and data normalized to infection rates in absence of serum. Values represent means of 3 technical replicates ± sem. Inhibitory titers 50 (Titer_50_) were calculated by nonlinear regression. SARS-CoV-2-reactive immunoglobulins (Ig) A, M, and G were determined by ELISA or chemiluminescent immunoassay (CLIA), values represent determined optic densities (OD). Sera are considered positive at ODs ≥ 1.1 or ≥ 1.0 in ELISA or CLIA, respectively. N/A not available.

## Discussion

We here describe a novel assay that allows quantification SARS-CoV-2 infection by measuring intracellular levels of the viral S protein in bulk cell cultures. The assay is based on the detection of *de novo* synthesized S protein by a S2-targeting antibody, and quantification via a corresponding secondary horseradish peroxidase (HRP)-linked antibody. This more sensitively detects nuances of viral replication than counting infected cells and is faster than determining titers of progeny virus. At high viral input (e.g. MOI 3), infection can already be detected after 24 hours, and at low viral input (e.g. MOI 0.005) after 48 hours. The assay has a linear range and signal-to-noise (S/N) ratios that are suitable to accurately determine the antiviral activity of drugs (IC_50_s), as shown for entry blocker EK1 (Xia et al., 2020a, 2020b), or intracellularly acting inhibitors remdesivir and lopinavir (Jeon et al., 2020; M. Wang et al., 2020). Additionally, the assay can be applied to detect and quantify titers of neutralizing sera of COVID-19 patients, all within only 2 days. This in-cell ELISA is easy to perform and follows standard ELISA readouts using HRP-mediated TMB substrate conversion and OD measurements after acidification with no need for expensive equipment.

Notably, the assay has been developed to be carried out in microtiter plates and should allow a convenient medium-to-high throughput testing of antivirals, antibodies, or antisera with timely availability of results, which is in the fast development of antivirals and in diagnostics. Due to targeting a highly conserved region and the relatively high sequence homology of global SARS-CoV-2 isolates, it is also applicable to other isolates as those that were tested herein. Furthermore, conservation in between related viruses suggest that, also SARS-CoV, and related civet SARS-CoV and bat SARS-like coronavirus infection can be detected with this assay (Ng et al., 2014; Walls et al., 2020). The in-cell ELISA was established using permissive Vero E6 and Caco-2 cells and confirmed using Calu-3 cells, but principally all other cell lines or primary cells supporting productive SARS-CoV-2 infection may also be used. In addition, SARS-CoV-2 is a BSL-3 pathogen which requires high safety requirements, which are usually at the expense of throughput. One additional advantage of the in-cell ELISA is that treatment of cells with paraformaldehyde results in the fixation and inactivation of virions, allowing a downstream processing of the plates outside a BSL-3 facility.

Another application of the in-cell S protein ELISA is to reliably determine infectious viral titres in virus stocks, cell culture supernatants or from patient swabs. Viral titers are usually quantified by limiting dilution analysis and microscopic determination of infected wells or staining of SARS-CoV-2 induced plaques or foci with crystal violet, neutral red or specific antibodies for SARS-CoV-2 antigens. We found that the in-cell ELISA allows to i) discriminate infected from uninfected wells, and ii) even after infection with very low MOIs (as low as 0.000005, which corresponds to one virion per three wells) at 4 days post infection (Fig. 3), representing an alternative for non-biased determining the TCID_50_ without the need of counting infected wells or plaques.

Conclusively, the S protein specific in-cell ELISA quantifies SARS-CoV-2 infection rates of different cell lines and allows to rapidly screen for and determine the potency of antiviral compounds. Thus, it represents a promising, rapid, readily available and easy to implement alternative to the current repertoire of laboratory techniques studying SARS-CoV-2 and will facilitate future research and drug development on COVID-19.

## Data sharing

Raw data is available upon request.

## Acknowledgments

We thank Daniela Krnavek, Nicola Schrott, Merve Karacan, Carolin Ludwig, Tirza Braun and Vivien Prex for experimental assistance. This project has received funding from the European Union’s Horizon 2020 research and innovation programme under grant agreement No 101003555 (Fight-nCoV) to J.M., the German Research Foundation (CRC1279) to J.M., S.S. and K.M.J.S., and an individual research grant (to J.A.M.). J.A.M. is indebted to the Baden-Württemberg Stiftung for the financial support of this research project by the Eliteprogramme for Postdocs. C.C., R.G., and D.S. are part of and R.G. is funded by a scholarship from the International Graduate School in Molecular Medicine Ulm. H.S. and B.J. receive funding from the German Ministry of Health for a clinical trial of convalescent plasma to treat severe COVID-19.

## References

Andreas, N.J., Kampmann, B., Mehring Le-Doare, K., 2015. Human breast milk: A review on its composition and bioactivity. Early Hum. Dev. 91, 629–635. https://doi.org/10.1016/j.earlhumdev.2015.08.013

Aubry, M., Richard, V., Green, J., Broult, J., Musso, D., 2016. Inactivation of Zika virus in plasma with amotosalen and ultraviolet A illumination. Transfusion 56, 33–40. https://doi.org/10.1111/trf.13271

Chanan, R.C., Shahani, K.M., Holly, R.G., 1964. Lysozyme Content of Human Milk. Nature 204, 76–77. https://doi.org/10.1038/204076a0

Chin, A.W.H., Chu, J.T.S., Perera, M.R.A., Hui, K.P.Y., Yen, H., Chan, M.C.W., Peiris, M., Poon, L.L.M., 2020. Stability of SARS-CoV-2 in different environmental conditions. The Lancet Microbe 1. e10. https://doi.org/10.1016/S2666-5247(20)30003-3

Chu, H., Chan, J.F.-W., Yuen, T.T.-T., Shuai, H., Yuan, S., Wang, Y., Hu, B., Yip, C.C., Tsang, J.O.-L., Huang, X., Chai, Y., Yang, D., Hou, Y., Chik, K.K.-H., Zhang, X., Fung, A.Y.-F., Tsoi, H.-W., Cai, J., Chan, W.-M., Ip, J.D., Chu, A.W., Zhou, J., Lung, D.C., Kok, K., To, K.K., Tsang, O.T., Chan, K., Yuen, K., 2020. Comparative tropism, replication kinetics, and cell damage profiling of SARS-CoV-2 and SARS-CoV with implications for clinical manifestations, transmissibility, and laboratory studies of COVID-19: an observational study. The Lancet Microbe 1, e14–e23. https://doi.org/10.1016/S2666-5247(20)30004-5

Conzelmann, Zou, Groß, Harms, Röcker, Riedel, Münch, Müller, 2019. Storage-Dependent Generation of Potent Anti-ZIKV Activity in Human Breast Milk. Viruses 11, 591. https://doi.org/10.3390/v11070591

Dajani, R., Zhang, Y., Taft, P.J., Travis, S.M., Starner, T.D., Olsen, A., Zabner, J., Welsh, M.J., Engelhardt, J.F., 2005. Lysozyme Secretion by Submucosal Glands Protects the Airway from Bacterial Infection. Am. J. Respir. Cell Mol. Biol. 32, 548–552. https://doi.org/10.1165/rcmb.2005-0059OC

Hadfield, J., Megill, C., Bell, S.M., Huddleston, J., Potter, B., Callender, C., Sagulenko, P., Bedford, T., Neher, R.A., 2018. NextStrain: Real-time tracking of pathogen evolution. Bioinformatics 34, 4121–4123. https://doi.org/10.1093/bioinformatics/bty407

Hoffmann, M., Kleine-Weber, H., Schroeder, S., Krüger, N., Herrler, T., Erichsen, S., Schiergens, T.S., Herrler, G., Wu, N.-H., Nitsche, A., Müller, M.A., Drosten, C., Pöhlmann, S., 2020. SARS-CoV-2 Cell Entry Depends on ACE2 and TMPRSS2 and Is Blocked by a Clinically Proven Protease Inhibitor. Cell 181, 271–280.e8. https://doi.org/10.1016/j.cell.2020.02.052

Jeon, S., Ko, M., Lee, J., Choi, I., Byun, S.Y., Park, S., Shum, D., Kim, S., 2020. Identification of antiviral drug candidates against SARS-CoV-2 from FDA-approved drugs. Antimicrob. Agents Chemother. 94. https://doi.org/10.1128/AAC.00819-20

Keil, S.D., Ragan, I., Yonemura, S., Hartson, L., Dart, N.K., Bowen, R., 2020. Inactivation of severe acute respiratory syndrome coronavirus 2 in plasma and platelet products using a riboflavin and ultraviolet light-based photochemical treatment. Vox Sang. vox.12937. https://doi.org/10.1111/vox.12937

Koenig, Á., Diniz, E.M. de A., Barbosa, S.F.C., Vaz, F.A.C., 2005. Immunologic Factors in Human Milk: The Effects of Gestational Age and Pasteurization. J. Hum. Lact. 21, 439–443. https://doi.org/10.1177/0890334405280652

Liu, J., Cao, R., Xu, M., Wang, X., Zhang, H., Hu, H., Li, Y., Hu, Z., Zhong, W., Wang, M., 2020. Hydroxychloroquine, a less toxic derivative of chloroquine, is effective in inhibiting SARS-CoV-2 infection in vitro. Cell Discov. 6, 16. https://doi.org/10.1038/s41421-020-0156-0

Liu, X., Li, Z., Liu, S., Sun, J., Chen, Z., Jiang, M., Zhang, Q., Wei, Y., Wang, X., Huang, Y.-Y., Shi, Y., Xu, Y., Xian, H., Bai, F., Ou, C., Xiong, B., Lew, A.M., Cui, J., Fang, R., Huang, H., Zhao, J., Hong, X., Zhang, Y., Zhou, F., Luo, H.-B., 2020. Potential therapeutic effects of dipyridamole in the severely ill patients with COVID-19. Acta Pharm. Sin. B 1–11. https://doi.org/10.1016/j.apsb.2020.04.008

Ma, Q., Pan, W., Li, R., Liu, B., Li, C., Xie, Y., Wang, Z., Zhao, J., Jiang, H., Huang, J., Shi, Y., Dai, J., Zheng, K., Li, X., Yang, Z., 2020. Liu Shen capsule shows antiviral and anti-inflammatory abilities against novel coronavirus SARS-CoV-2 via suppression of NF-κB signaling pathway. Pharmacol. Res. 104850. https://doi.org/10.1016/j.phrs.2020.104850

Manenti, A., Maggetti, M., Casa, E., Martinuzzi, D., Torelli, A., Trombetta, C.M., Marchi, S., Montomoli, E., 2020. Evaluation of SARS-CoV-2 neutralizing antibodies using a CPE-based colorimetric live virus micro-neutralization assay in human serum samples. J. Med. Virol. jmv. 25986. https://doi.org/10.1002/jmv.25986

Mauthe, M., Orhon, I., Rocchi, C., Zhou, X., Luhr, M., Hijlkema, K.-J., Coppes, R.P., Engedal, N., Mari, M., Reggiori, F., 2018. Chloroquine inhibits autophagic flux by decreasing autophagosomelysosome fusion. Autophagy 14, 1435–1455. https://doi.org/10.1080/15548627.2018.1474314

McDermott, A.M., 2013. Antimicrobial compounds in tears. Exp. Eye Res. 117, 53–61. https://doi.org/10.1016/j.exer.2013.07.014

Monteil, V., Kwon, H., Prado, P., Hagelkrüys, A., Wimmer, R.A., Stahl, M., Leopoldi, A., Garreta, E., Hurtado del Pozo, C., Prosper, F., Romero, J.P., Wirnsberger, G., Zhang, H., Slutsky, A.S., Conder, R., Montserrat, N., Mirazimi, A., Penninger, J.M., 2020. Inhibition of SARS-CoV-2 Infections in Engineered Human Tissues Using Clinical-Grade Soluble Human ACE2. Cell 181, 905–913.e7. https://doi.org/10.1016/j.cell.2020.04.004

Müller, J.A., Harms, M., Krüger, F., Groß, R., Joas, S., Hayn, M., Dietz, A.N., Lippold, S., von Einem, J., Schubert, A., Michel, M., Mayer, B., Cortese, M., Jang, K.S., Sandi-Monroy, N., Deniz, M., Ebner, F., Vapalahti, O., Otto, M., Bartenschlager, R., Herbeuval, J.-P., Schmidt-Chanasit, J., Roan, N.R., Münch, J., 2018. Semen inhibits Zika virus infection of cells and tissues from the anogenital region. Nat. Commun. 9, 2207. https://doi.org/10.1038/s41467-018-04442-y

Müller, J.A., Harms, M., Schubert, A., Mayer, B., Jansen, S., Herbeuval, J.-P.J.-P., Michel, D., Mertens, T., Vapalahti, O., Schmidt-Chanasit, J., Münch, J., 2017. Development of a high-throughput colorimetric Zika virus infection assay. Med. Microbiol. Immunol. 206, 175–185. https://doi.org/10.1007/s00430-017-0493-2

Ng, O.W., Keng, C.T., Leung, C.S.W., Peiris, J.S.M., Poon, L.L.M., Tan, Y.J., 2014. Substitution at aspartic acid 1128 in the SARS coronavirus spike glycoprotein mediates escape from a S2 domain-targeting neutralizing monoclonal antibody. PLoS One 9, 1–11. https://doi.org/10.1371/journal.pone.0102415

Ou, X., Liu, Y., Lei, X., Li, P., Mi, D., Ren, L., Guo, L., Guo, R., Chen, T., Hu, J., Xiang, Z., Mu, Z., Chen, X., Chen, J., Hu, K., Jin, Q., Wang, J., Qian, Z., 2020. Characterization of spike glycoprotein of SARS-CoV-2 on virus entry and its immune cross-reactivity with SARS-CoV. Nat. Commun. 11, 1620. https://doi.org/10.1038/s41467-020-15562-9

Petit, J.F., Jollès, P., 1963. Purification and Analysis of Human Saliva Lysozyme. Nature 200, 168–169. https://doi.org/10.1038/200168a0

Röcker, A.E., Müller, J.A., Dietzel, E., Harms, M., Krüger, F., Heid, C., Sowislok, A., Riber, C.F., Kupke, A., Lippold, S., von Einem, J., Beer, J., Knöll, B., Becker, S., Schmidt-Chanasit, J., Otto, M., Vapalahti, O., Zelikin, A.N., Bitan, G., Schrader, T., Münch, J., 2018. The molecular tweezer CLR01 inhibits Ebola and Zika virus infection. Antiviral Res. 152, 26–35. https://doi.org/10.1016/j.antiviral.2018.02.003

Runfeng, L., Yunlong, H., Jicheng, H., Weiqi, P., Qinhai, M., Yongxia, S., Chufang, L., Jin, Z., Zhenhua, J., Haiming, J., Kui, Z., Shuxiang, H., Jun, D., Xiaobo, L., Xiaotao, H., Lin, W., Nanshan, Z., Zifeng, Y., 2020. Lianhuaqingwen exerts anti-viral and anti-inflammatory activity against novel coronavirus (SARS-CoV-2). Pharmacol. Res. 156, 104761. https://doi.org/10.1016/j.phrs.2020.104761

Walls, A.C., Park, Y.J., Tortorici, M.A., Wall, A., McGuire, A.T., Veesler, D., 2020. Structure, Function, and Antigenicity of the SARS-CoV-2 Spike Glycoprotein. Cell 181, 281–292.e6. https://doi.org/10.1016/j.cell.2020.02.058

Wang, C., Li, W., Drabek, D., Okba, N.M.A., Haperen, R. Van, Osterhaus, A.D.M.E., Kuppeveld, F.J.M. Van, Haagmans, B.L., Grosveld, F., Bosch, B., 2020. A human monoclonal antibody blocking SARS-CoV-2 infection. Nat. Commun. 11, 1–6. https://doi.org/10.1038/s41467-020-16256-y

Wang, M., Cao, R., Zhang, L., Yang, X., Liu, J., Xu, M., Shi, Z., Hu, Z., Zhong, W., Xiao, G., 2020. Remdesivir and chloroquine effectively inhibit the recently emerged novel coronavirus (2019-nCoV) in vitro. Cell Res. 30, 269–271. https://doi.org/10.1038/s41422-020-0282-0

Wang, Q., Zhang, Y., Wu, L., 2020. Structural and Functional Basis of SARS-CoV-2 Entry by Using Human ACE2. Cell 181, 894–904. https://doi.org/10.1016/j.cell.2020.03.045

Xia, S., Liu, M., Wang, C., Xu, W., Lan, Q., Feng, S., Qi, F., Bao, L., Du, L., Liu, S., Qin, C., Sun, F., Shi, Z., Zhu, Y., Jiang, S., Lu, L., 2020a. Inhibition of SARS-CoV-2 (previously 2019-nCoV) infection by a highly potent pan-coronavirus fusion inhibitor targeting its spike protein that harbors a high capacity to mediate membrane fusion. Cell Res. 2. https://doi.org/10.1038/s41422-020-0305-x

Xia, S., Zhu, Y., Liu, M., Lan, Q., Xu, W., Wu, Y., Ying, T., Liu, S., Shi, Z., Jiang, S., Lu, L., 2020b. Fusion mechanism of 2019-nCoV and fusion inhibitors targeting HR1 domain in spike protein. Cell. Mol. Immunol. 3–5. https://doi.org/10.1038/s41423-020-0374-2

Yao, X., Ye, F., Zhang, M., Cui, C., Huang, B., Niu, P., Liu, X., Zhao, L., Dong, E., Song, C., Zhan, S., Lu, R., Li, H., Tan, W., Liu, D., 2020. In Vitro Antiviral Activity and Projection of Optimized Dosing Design of Hydroxychloroquine for the Treatment of Severe Acute Respiratory Syndrome Coronavirus 2 (SARS-CoV-2). Clin. Infect. Dis. 1–8. https://doi.org/10.1093/cid/ciaa237

Zhu, N., Zhang, D., Wang, W., Li, X., Yang, B., Song, J., Zhao, X., Huang, B., Shi, W., Lu, R., Niu, P., Zhan, F., Ma, X., Wang, D., Xu, W., Wu, G., Gao, G.F., Tan, W., 2020. A Novel Coronavirus from Patients with Pneumonia in China, 2019. N. Engl. J. Med. 382, 727–733. https://doi.org/10.1056/NEJMoa2001017

